# Interleukin-4 suppresses TIMP-2 transcription during endometriosis: Allele-specific regulation of the TIMP-2 −418G/C promoter by SP1

**DOI:** 10.64898/2026.10.15.759364

**Authors:** Susmita Saha, Supratim Ghosh, Tapasi Roy, Debasis Nayak, Pramathes Das Mahapatra, Arindam Maitra, Snehasikta Swarnakar

**Affiliations:** Division of Infectious Diseases & Immunology, CSIR-Indian Institute of Chemical Biology, Kolkata, West Bengal, India; Biotechnology Research and Innovation Council -National Institute of Biomedical Genomics (BRIC-NIBMG), Kalyani, West Bengal, India; Department of Obstetrics and Gynecology, Spectrum Clinic and Endoscopy Research Institute, Kolkata, West Bengal, India; Division of Cancer Biology & Inflammatory Disorder, CSIR-Indian Institute of Chemical Biology, Kolkata, West Bengal, India

**Keywords:** – Endometriosis, TIMP-2, IL-4 signaling, SP1, ECM, Promoter polymorphism, Cell type, Cell state

## Abstract

Tissue inhibitor of metalloproteinase-2 (TIMP-2) is a critical regulator of extracellular matrix (ECM) homeostasis in inflammatory disorder, but the mechanisms underlying its dysregulation in endometriosis remain poorly understood. Furthermore, whether inflammatory cytokine signaling and functional genetic variants within the TIMP-2 promoter contribute to its altered expression during disease progression is unknown. This study investigated the regulation of TIMP-2 expression by interleukin-4 (IL-4) signaling and the functional significance of the TIMP-2 −418G/C promoter polymorphism in endometriosis. We found that TIMP-2 expression was significantly reduced in ectopic lesions and serum samples from endometriosis patients in comparison to normal. Functional promoter analyses revealed that the −418C allele exhibited higher basal transcriptional activity than the G allele while displayed greater sensitivity to IL-4-mediated repression. IL-4 activated STAT6 signaling concomitant with SP1 and TIMP-2 downregulation, whereas SP1 overexpression restored TIMP-2 expression and SP1 silencing further suppressed its transcription, identifying SP1 as a positive regulator of TIMP-2. Single-cell transcriptomic analyses identified fibroblasts and mesenchymal populations as the major TIMP-2-expressing cell types and revealed enhanced inflammatory signaling and ECM remodeling networks in endometriosis. Collectively, these findings identify an IL-4–SP1– TIMP-2 regulatory axis that contributes to aberrant ECM turnover in endometriosis. Furthermore, the TIMP-2 −418G/C promoter polymorphism influences cytokine-mediated transcriptional regulation of TIMP-2, providing new mechanistic insights into the interplay between inflammatory signaling, genetic variation, and ECM homeostasis during endometriosis progression.

## Introduction

Endometriosis is a chronic inflammatory gynecological disorder affecting approximately 10– 15% of women of reproductive age and is a major cause of chronic pelvic pain, infertility, and reduced quality of life [1]. The disease is characterized by the presence of endometrial-like tissue outside the uterine cavity, most commonly on the ovaries, peritoneum, and pelvic ligaments. Although its etiology remains incompletely understood, endometriosis is widely recognized as a multifactorial disease involving aberrant hormonal signaling, immune dysregulation, chronic inflammation, and extracellular matrix (ECM) remodeling [2]. A defining feature of endometriosis is the ability of ectopic endometrial cells to adhere, invade, and establish persistent lesions at distant anatomical sites [3] [4]. These processes are closely associated with excessive ECM remodeling, fibrosis, and adhesion formation, which contribute to pelvic pain, infertility, and progressive distortion of pelvic anatomy [5] [6]. The establishment and progression of endometriotic lesions are strongly influenced by dysregulated ECM turnover and persistent inflammatory signaling. Cytokines such as transforming growth factor-β1 (TGF-β1) and interleukin-4 (IL-4) have been implicated in fibrosis and tissue remodeling within the endometriotic microenvironment [7] [8]. TGF-β1 promotes fibroblast-to-myofibroblast differentiation and excessive collagen deposition, whereas IL-4 contributes to fibroblast proliferation and ECM accumulation through the induction of collagen-associated genes and connective tissue growth factors [9] [10]. Elevated levels of IL-4 in the peritoneal fluid and endometriotic lesions suggest that persistant cytokine signaling contributes to aberrant tissue repair and lesion formation during endometriosis [11] [12] [13]. Matrix metalloproteinases (MMPs) and their endogenous tissue inhibitors (TIMPs) are central regulators of ECM homeostasis. Among them, tissue inhibitor of metalloproteinase-2 (TIMP-2) plays a critical role in maintaining ECM integrity by regulating MMP activity and controlling matrix degradation [14]. In addition to its inhibitory effects on proteolysis, TIMP-2 also influences cell–matrix interactions, angiogenesis, and tissue remodeling [15], [16]. Altered TIMP-2 expression has been associated with enhanced invasion, fibrosis, and abnormal tissue remodeling in endometriosis [17]. Previous studies have further suggested that TIMP-2 expression may be influenced by inflammatory cytokines and genetic polymorphisms within its promoter sequence [18], [19], [20]. However, the molecular mechanisms regulating TIMP-2 transcription under inflammatory conditions in endometriosis remain poorly understood. A functional polymorphism within the TIMP-2 promoter region, −418 G>C, has been reported to influence transcriptional activity and disease susceptibility. This polymorphic substitution lies within a predicted SP1-binding site and influence basal promoter activity as well as transcription factor recruitment [21],[22]. SP1 is an important transcription factor involved in the regulation of genes associated with ECM remodeling, inflammation, and cellular proliferation. Previous studies have demonstrated that SP1 and NF-Y cooperatively regulate TIMP-2 promoter activity, suggesting that alterations at the −418 site may influence TIMP-2 expression in a context-dependent manner [23], [24]. Clinically, the −418 G/C polymorphism has been associated with altered susceptibility to several pathological conditions, including endometriosis and cancer [20]. In addition to genetic regulation, inflammatory cytokine signaling may further modulate TIMP-2 transcriptional activity. In support IL-4 has been reported to be elevated in endometriosis and is known to regulate multiple transcriptional pathways associated with fibrosis and chronic inflammation [25], [26], [27]. Evidence suggesting a correlation between IL-4 signaling and TIMP-2 expression in adhesion-associated tissues raises the possibility that inflammatory cytokines may regulate TIMP-2 through SP1-dependent transcriptional mechanisms [28]. In the present study, we investigated the molecular mechanisms regulating TIMP-2 expression in endometriosis through genetic, inflammatory, and transcriptional analyses. We further examined the association of the TIMP-2 −418 G/C promoter polymorphism with endometriosis susceptibility in an eastern Indian population and evaluated the functional activity of the G and C allelic promoter variants under basal, IL-4– stimulated, and SP1-modulated conditions. In parallel, tissue-based analyses and single-cell transcriptomic profiling were performed to characterize TIMP-2 expression profile, inflammatory signaling pathways, and cell–cell communication networks associated with endometriosis progression.

## 2. Materials and Methods

### 2.1. Clinical samples and ethics approval

Patients presenting with symptoms including primary or secondary infertility, abdominal pain, rectal pain, and chest pain were referred to the Gynaecology Division of Spectrum Clinic & Endoscopy Research Institute (Kolkata, India), where laparoscopic evaluation was performed for diagnostic confirmation of endometriosis. Endometriotic lesions were staged according to the revised American Society for Reproductive Medicine (rASRM) classification system.

The study protocol was approved by the Human Ethics Committee of the Indian Institute of Chemical Biology (Approval No. IICB/IRB/SS02-2015) and the Human Ethics Committee of Spectrum Clinic & Endoscopy Research Institute (Approval No. SCERI-IEC015). Written informed consent was obtained from all participants prior to sample collection.

The study cohort represented the eastern Indian population, including West Bengal and neighboring states. Women aged 18–40 years with surgically confirmed endometriosis were included in the study. Exclusion criteria included pregnancy, lactation, previous hysterectomy or bilateral oophorectomy, and the presence of other gynecological or oncological disorders. Clinical history, demographic information, and disease characteristics were recorded for all participants. Tissue grading and blood collection were performed during the proliferative phase of the menstrual cycle. Control samples were obtained from women with proven fertility (≥1 child) and no clinical or laparoscopic evidence of endometriosis or other gynecological diseases.

### 2.2. Serum and tissue collection

Peripheral blood samples were collected aseptically in EDTA-coated tubes and stored at 4 °C until genomic DNA extraction using the QIAamp DNA Blood Midi Kit (Qiagen, Germany). Serum samples were separated, supplemented with protease inhibitor cocktail (Sigma-Aldrich, USA), and stored at −80 °C until further analysis.

Tissue samples were divided into two portions immediately after collection. One portion was fixed in 4% paraformaldehyde for immunofluorescence studies, while the remaining tissue was snap-frozen and stored at −80 °C for molecular and protein analyses.

### 2.3. Genotyping of TIMP-2 −418 G/C polymorphism

Genotyping of the TIMP-2 −418 G/C polymorphism was performed using the polymerase chain reaction–restriction fragment length polymorphism (PCR–RFLP) method. The promoter region was amplified using the following primers:

Forward: 5′-CGTCTCTTGTTGGCTGGTCA-3′
Reverse: 5′-CCTTCAGCTCGACTCTGGAG-3′

Each PCR reaction contained 10 ng genomic DNA, 0.3 mM of each primer, and 10 μL Premix Taq V2.0 (XcelGen, India). Amplification was carried out under the following conditions: initial denaturation at 95 °C for 5 min, followed by 30 cycles of denaturation at 95 °C for 30 s, annealing at 55 °C for 30 s, and extension at 72 °C for 30 s, with a final extension step at 72 °C for 7 min. Amplified products were digested with BsoBI restriction enzyme (Takara Bio, Japan) and resolved on 2% agarose gels containing ethidium bromide.

### 2.4. Generation of TIMP-2 promoter–reporter constructs

The TIMP-2 promoter region was amplified from genomic DNA obtained from individuals carrying either the GG or CC genotype at the −418 site. The following primers were used:

Forward: 5′-GAGGTACCGGCCACCCGCCCACAGAAACGGGG-3′
Reverse: 5′-CAGAGCTCGCGGCCGCGCTGCCTTCTACGGATGTGTTTG-3′

KpnI and SacI restriction sites are indicated within the primer sequences. Amplified promoter fragments spanning the −499 to +32 bp region of the TIMP-2 promoter were cloned into the pGL4.72 luciferase reporter vector (Promega, USA) at the KpnI and SacI restriction sites.

Constructs containing either the G or C allele at the −418 polymorphic site were confirmed by DNA sequencing. Endotoxin-free plasmid DNA was prepared using the Promega Endotoxin-Free Plasmid DNA Purification Kit.

### 2.5. Cell culture

Human endometrial epithelial cell lines Ishikawa and RL95-2, endometrial stromal cell line KC02-44D, ovarian adenocarcinoma cell lines SK-OV-3 and OVCAR-3, and breast adenocarcinoma cell line MCF-7 were obtained from the American Type Culture Collection (ATCC, USA). Ishikawa, RL95-2, and KC02-44D cells were maintained in Dulbecco’s Modified Eagle Medium/F12 (DMEM/F12; Gibco, USA) without phenol red. MCF-7 cells were cultured in Dulbecco’s Modified Eagle Medium (DMEM; Gibco), whereas SK-OV-3 and OVCAR-3 cells were maintained in RPMI-1640 medium (Gibco). OVCAR-3 cells were supplemented with 20% fetal bovine serum (FBS). All media were supplemented with 10% FBS, 100 U/mL penicillin, and 100 μg/mL streptomycin. Cells were maintained at 37 °C in a humidified atmosphere containing 5% CO₂.

### 2.6. Luciferase reporter assays and IL-4 stimulation

Transient transfection assays were performed using Lipofectamine 2000 (Invitrogen, USA) according to the manufacturer’s instructions. Cells were seeded at approximately 60% confluence and transfected with 1 μg of either G-or C-allele TIMP-2 promoter–reporter constructs.

For cytokine stimulation experiments, recombinant human IL-4 (R&D Systems, USA) was added at concentrations of 1, 2.5, 5, 7.5, and 10 ng/mL for 12 h. In selected experiments, Ishikawa cells were transfected with promoter constructs and treated with IL-4 (10 ng/mL) 6 h post-transfection, followed by an additional 12 h incubation. Luciferase activity was measured using a luminometer and normalized to total protein concentration determined by Bradford assay. Results were expressed as relative light units (RLU). All transfection experiments were independently performed at least three times.

### 2.7. SP1 overexpression and siRNA-mediated knockdown assays

For SP1 overexpression studies, Ishikawa cells were co-transfected with TIMP-2 promoter– reporter constructs (1 μg/well) and increasing concentrations of SP1 expression plasmid (100, 250, 500 ng, and 1 μg). Empty pcDNA3.1 vector was used as a balancing control. For RNA interference experiments, Ishikawa cells were co-transfected with G- or C-allele promoter constructs and SP1-specific siRNA (10, 20, or 30 nM; Santa Cruz Biotechnology, USA) using Lipofectamine RNAiMAX (Invitrogen). Scrambled siRNA was used as a negative control. Luciferase activity was analyzed 12 h following IL-4 treatment or siRNA transfection and normalized to total protein concentration.

### 2.8. Protein extraction and immunoblot analysis

Cells and tissue samples were homogenized in lysis buffer containing 10 mM Tris-HCl (pH 8.0), 150 mM NaCl, 1% Triton X-100, and protease inhibitor cocktail (Sigma-Aldrich). Homogenates were centrifuged at 12,000 × g for 15 min at 4 °C, and supernatants were collected for protein analysis. Protein concentrations were determined using Bradford and Lowry assays. Equal amounts of protein (70–100 μg) were resolved on 10% SDS-PAGE gels and transferred onto nitrocellulose membranes. Membranes were incubated with primary antibodies against TIMP-2 (Cell Signaling Technology, #5738), SP1 (Cell Signaling Technology, #9389), phospho-SP1 (Thr453; Abcam, ab59257), GAPDH (Santa Cruz Biotechnology, sc-9104), and β-actin (Cell Signaling Technology, #4967). Appropriate HRP-conjugated secondary antibodies (Millipore, USA) were subsequently applied, and immunoreactive bands were visualized using Immobilon HRP substrate (Millipore). For time-course experiments, Ishikawa cells were treated with recombinant IL-4 (10 ng/mL) for 10, 30, 60, 120, 180, 240, 360, and 720 min prior to protein extraction.

### 2.9. Enzyme-linked immunosorbent assay (ELISA)

TIMP-2 and IL-4 concentrations in serum and tissue lysates were quantified using commercially available human TIMP-2 and IL-4 ELISA kits (R&D Systems, USA) according to the manufacturer’s instructions.

### 2.10. Immunofluorescence analysis

For tissue immunofluorescence, biopsy samples were fixed in 4% paraformaldehyde, dehydrated through graded ethanol series, and embedded in paraffin. Sections (5 μm) were deparaffinized in xylene and rehydrated through decreasing alcohol concentrations. Antigen retrieval was performed using 0.05% trypsin containing 0.1% CaCl₂, followed by blocking with 5% bovine serum albumin (BSA). Sections were incubated overnight at 4 °C with primary antibodies diluted in TBS containing 1% BSA. Following washing with TBST, samples were incubated with FITC-conjugated secondary antibodies (Santa Cruz Biotechnology) and counterstained with propidium iodide. For cellular immunofluorescence assays, Ishikawa cells were seeded in poly-L-lysine–coated 24-well plates and treated with recombinant IL-4 (10 ng/mL) for 12 h. Cells were fixed in 4% paraformaldehyde, permeabilized with 0.1% Triton X-100, and blocked with 1% BSA in TBST. Cells were incubated overnight at 4 °C with primary antibodies, followed by FITC- and Texas Red–conjugated secondary antibodies (Santa Cruz Biotechnology). Nuclei were counterstained using Hoechst dye. Images were acquired using confocal microscopy at 10× and 40× magnifications.

### 2.11. Single-cell RNA sequencing (scRNA-seq) analysis

Single-cell RNA sequencing datasets comprising 48 samples were obtained from the Gene Expression Omnibus (GEO) under accession number GSE213216 [29]. For each sample, the cells with expression of less than 200 genes were removed to get rid of low-quality cells and empty droplets. Cells with more than 20% mitochondrial reads were filtered out to remove lysed cells and retain only good-quality cells using the R package Seurat v4.4.0 (https://github.com/satijalab/seurat) [30]. Subsequently, doublets were excluded @ 0.8% per 1000 cells using the R package DoubletFinder v2.0.3 (https://github.com/chris-mcginnis-ucsf/DoubletFinder) [31]. Dimensional reduction and cell clustering were performed for each sample using the Louvain algorithm in Seurat [30]. All the samples were then merged and SCTransform (https://github.com/satijalab/sctransform) normalized for regressing out effects of read depth per cell, percentage of mitochondrial reads per cell, and technical variation among samples [32]. The data from all samples were integrated using the R package Harmony v1.2.3 (https://github.com/immunogenomics/harmony) and visualized using uniform manifold approximation and projection (UMAP) [33]. The marker genes for each cell cluster were identified using Seurat FindAllMarkers() which essentially utilizes the Wilcoxon rank-sum test and clusters were annotated to each cell type accordingly [30]. From the overall cell types identified, epithelial cells, immune cells, and fibroblasts were separately reclustered, and gene expression of interest was determined using DoHeatmap(), DotPlot(), and VlnPlot() in Seurat. The cell-cell communication analyses were conducted using the R package CellChat (https://github.com/sqjin/CellChat) with ‘CellChatDB.human’ as the ligand-receptor interaction database [34]. For each cell type, the cell-cell interaction network at a single signaling pathway level was studied as well as incoming and outgoing communication networks were built for each cell type and compared across each disease condition.

### 2.12. Statistical analysis

Statistical analyses were performed using GraphPad InStat3 software (GraphPad Software Inc., USA). Data are presented as mean ± SEM unless otherwise stated. Differences between two groups were analyzed using Student’s t-test. Comparisons among multiple groups were performed using one-way ANOVA followed by Tukey’s post hoc test. Genotype distributions of the TIMP-2 −418 G/C polymorphism between endometriosis patients and healthy controls were analyzed using Fisher’s exact test. Odds ratios (ORs) and 95% confidence intervals (CIs) were calculated using binary logistic regression analysis, with the GG genotype used as the reference group. Associations between genotype and clinical parameters, including disease stage, pain, and lesion site, were evaluated using multivariate logistic regression analysis. All experiments were independently performed at least three times. Statistical significance was considered at p < 0.05.

### 2.13. Data availability

The single-cell RNA sequencing datasets analyzed in this study are publicly available in the Gene Expression Omnibus (GEO) repository under accession number GSE213216.

## 3. Results

### 3.1. TIMP-2 is consistently downregulated in endometrial tissues and serum of women with endometriosis vs without endometriosis

To confirm the reduced expression of TIMP-2 in an independent Eastern Indian cohort and establish the foundation for subsequent mechanistic investigations, TIMP-2 protein expression was examined in tissue and serum samples obtained from women with endometriosis and from patients coming to the clinic either with infertility-related problemsor with minor gynaecological problems. Western blot analysis demonstrated significantly lower TIMP-2 protein levels in ectopic endometriotic lesions compared with eutopic endometrial tissues from healthy controls (Fig. 1A). Similarly, eutopic endometrial tissues from women with endometriosis exhibited markedly reduced TIMP-2 expression relative to eutopic endometrium from disease-free controls (Fig. 1B). Consistent with these tissue-specific observations, serum samples from patients with endometriosis also showed significantly decreased TIMP-2 protein levels compared with healthy controls (Fig. 1C), indicating that reduced TIMP-2 expression is evident both locally and systemically.

**Figure 1.**
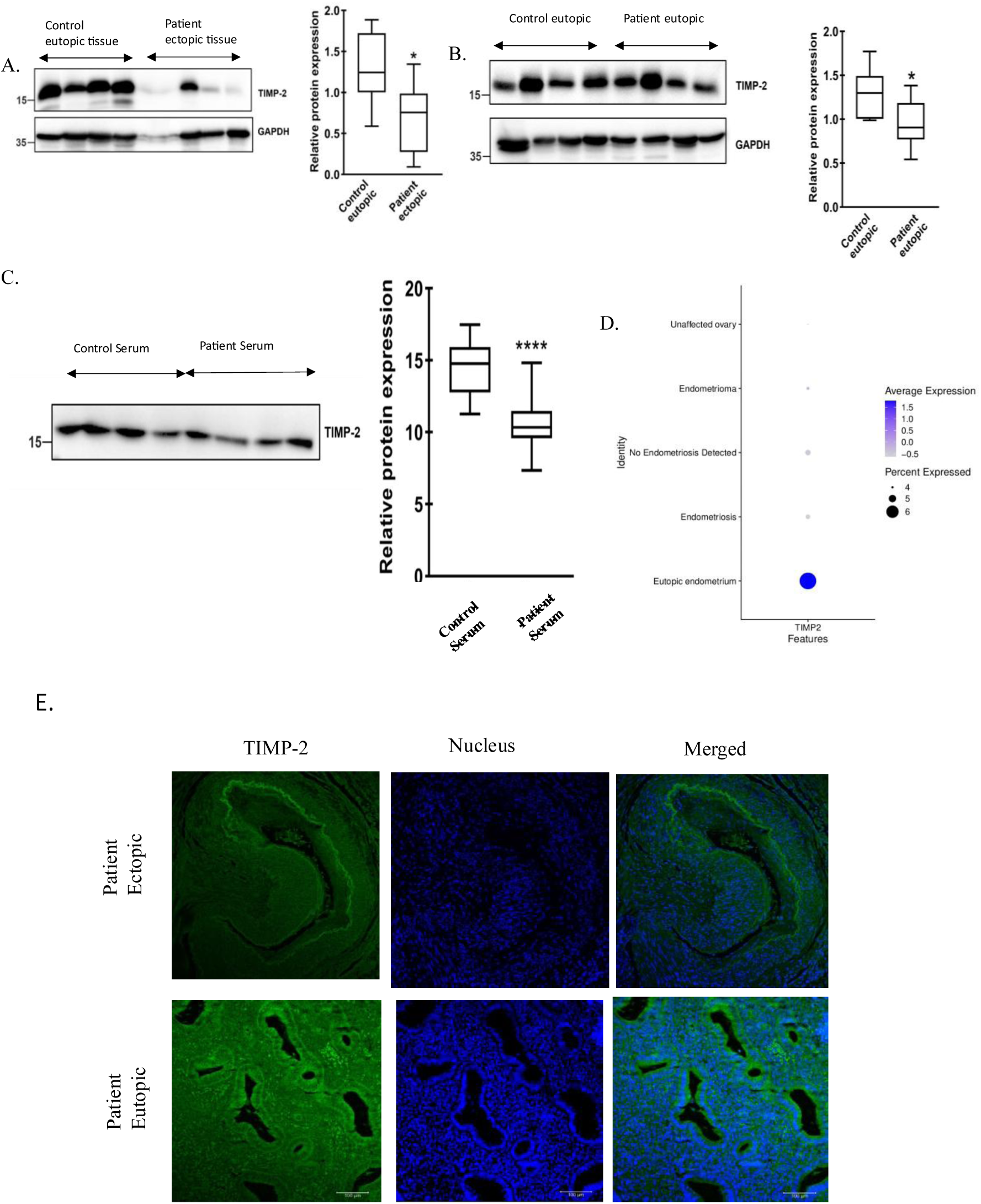
Reduced TIMP-2 expression in endometriotic tissues and serum from endometriosis patients (A) Representative images of immunoblot analysis of TIMP-2 expression in eutopic endometrium from healthy controls and ectopic lesions from endometriosis patients. Corresponding densitometric quantification normalized to GAPDH is shown on the right. (B) Immunoblot analysis comparing TIMP-2 protein expression between eutopic endometrium from healthy controls and eutopic endometrium from endometriosis patients. Densitometric quantification of TIMP-2 expression relative to GAPDH is presented alongside the blot. (C) Representative image of immunoblot showing TIMP-2 protein levels in serum samples obtained from healthy individuals and endometriosis patients. Quantitative analysis of serum TIMP-2 expression is shown on the right. (D) Dot plot showing *TIMP2* expression in different sample identities from the scRNA-seq dataset, including unaffected ovary, endometrioma, no endometriosis detected, endometriosis, and eutopic endometrium. Dot size indicates the percentage of cells expressing *TIMP2*, whereas color intensity represents the average expression level. (E) Representative immunofluorescence staining of TIM/P-2 in control eutopic endometrium, eutopic endometrium from endometriosis patients, and ectopic endometriotic lesions, demonstrating reduced TIMP-2 expression in disease tissues. Data are presented as mean ± SEM. Statistical significance was determined using Student’s t-test. *p < 0.05; ****p < 0.0001.

To further substantiate these findings at the transcriptomic level, we re-analyzed the publicly available single-cell RNA sequencing dataset (GSE213216) generated by Lawrenson and colleague. Dot plot analysis revealed that *TIMP2* expression was highest in eutopic endometrium from women without endometriosis, whereas its expression was markedly reduced in endometriosis-associated tissues, including endometriotic lesions and endometrioma (Fig. 1D), corroborating our protein expression data.

Immunofluorescence staining further confirmed the differential expression pattern of TIMP-2 within endometriosis tissues. Consistent with the western blot findings, TIMP-2 immunoreactivity was lower in ectopic lesions than in the corresponding eutopic endometrium from women with endometriosis (Fig. 1E), further supporting the progressive loss of TIMP-2 expression during lesion formation.

Collectively, these findings demonstrate that TIMP-2 downregulation is a typical molecular feature of endometriosis across tissue, serum, and independent transcriptomic datasets. Having established this disease-associated reduction in TIMP-2 expression, we next sought to identify the cellular distribution of *TIMP-2* and investigate the upstream regulatory mechanisms responsible for its suppression in endometriosis.

### 3.2. Single-cell transcriptomic analysis identifies the cellular distribution of *TIMP2* and implicates IL-4 signaling in endometriosis

Having established that TIMP-2 expression is reduced in endometriosis tissues and serum (Fig. 1), we next sought to determine the cellular distribution of *TIMP2* expression and identify potential upstream inflammatory regulators associated with its downregulation in specific disease-relevant cell types. Cell type-specific analysis revealed that *TIMP2* was predominantly expressed in fibroblasts, stromal cells, myofibroblasts and pericytes, whereas relatively low expression was detected in immune cell populations (Fig. 2A).

**Figure 2.**
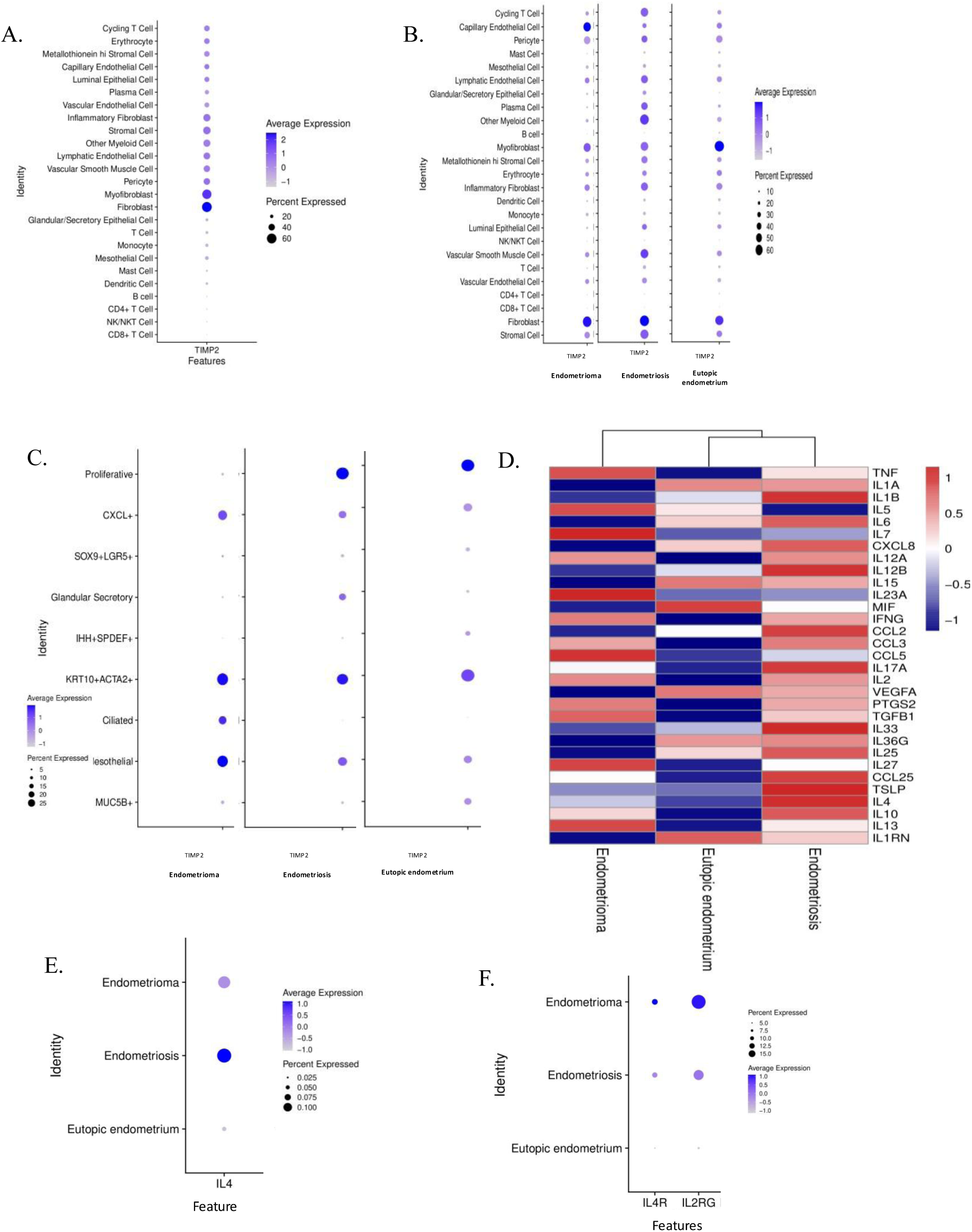
Single-cell transcriptomic analysis reveals differential *TIMP2* and IL-4 expression across endometrial tissue states. (A) Dot plot illustrating cell-type–specific *TIMP2* expression across major endometrial cell populations. (B) Distribution of *TIMP2* expression across individual cell populations within endometrioma, endometriosis, and eutopic endometrium. (C) Expression of *TIMP2*-associated epithelial cell-state markers across endometrial tissue conditions. (D) Heatmap showing differential expression of inflammatory cytokines, chemokines, and signaling-associated genes across tissue states. (E) Dot plot showing IL4 expression across endometrioma, endometriosis, and eutopic endometrium. (F) Dot plot showing expression of IL4 signaling-associated receptors including IL4R and IL2RG across endometrial tissue conditions. Dot size indicates the percentage of expressing cells and color intensity represents scaled expression levels.

We next investigated whether *TIMP2* expression within these cell populations was altered during disease progression. Comparative analysis demonstrated a marked reduction in TIMP2 expression in stromal cells, fibroblasts and other *TIMP2*-expressing populations in endometriosis and endometrioma compared with eutopic endometrium (Fig. 2B), indicating that the loss of *TIMP2* occurs across multiple stromal compartments rather than being restricted to a single cell type indicating system-level dysregulation of TIMP2 expression associated with endometrial pathogenesis.

Since epithelial cells constitute a major component of endometriotic lesions, *TIMP2* expression was further examined within epithelial subpopulations. Among these, the KRT10^+^ACTA2^+^ epithelial subset exhibited the highest *TIMP2* expression, whereas comparatively lower expression was observed in the remaining epithelial populations(Fig. 2C)[35].

To identify inflammatory mediators potentially associated with reduced *TIMP2* expression, cytokine expression profiles were subsequently analyzed. Most of the inflammatory cytokines were differentially upregulated in endometriosis followed by endometrioma compared to eutopic endometrium demonstrating systemic hyperinflammation in pathologic conditions (Fig. 2D). Among these, *IL4* expression was markedly increased in endometriosis and endometrioma compared with eutopic endometrium (Fig. 2E). Furthermore, the *IL4* receptor components, *IL4R* and *IL2RG*, also exhibited increased expression in endometriosis and endometrioma relative to eutopic endometrium (Fig. 2F), indicating enrichment of the IL-4 signaling machinery within the endometriotic microenvironment. Collectively, these cell type specific transcriptomic analyses demonstrate that *TIMP2* is progressively downregulated across multiple stromal and epithelial cell populations in endometriosis, while the concomitant enrichment of *IL4* together with its receptor components identifies the IL-4 signaling axis as a potential upstream regulator of TIMP2, providing the rationale for subsequent functional studies investigating the direct effect of IL-4 on TIMP-2 expression.

### 3.3. IL-4 suppresses TIMP-2 expression through activation of IL-4R/STAT6 signaling and modulation of SP1

Following the identification of IL-4 as a potential upstream regulator of TIMP2 from the single-cell transcriptomic analysis (Fig. 2), we next investigated whether IL-4 directly regulates TIMP-2 expression in endometrial epithelial cells. Ishikawa cells were treated with increasing concentrations of recombinant IL-4 (0–10 ng/ml) for 12 h, and TIMP-2 levels in the conditioned medium were analyzed by immunoblotting. IL-4 treatment resulted in a dose-dependent reduction in TIMP-2 expression, with significant suppression observed at 7.5 and 10 ng/ml (Fig. 3A), indicating that IL-4 negatively regulates TIMP-2 expression.

**Figure 3.**
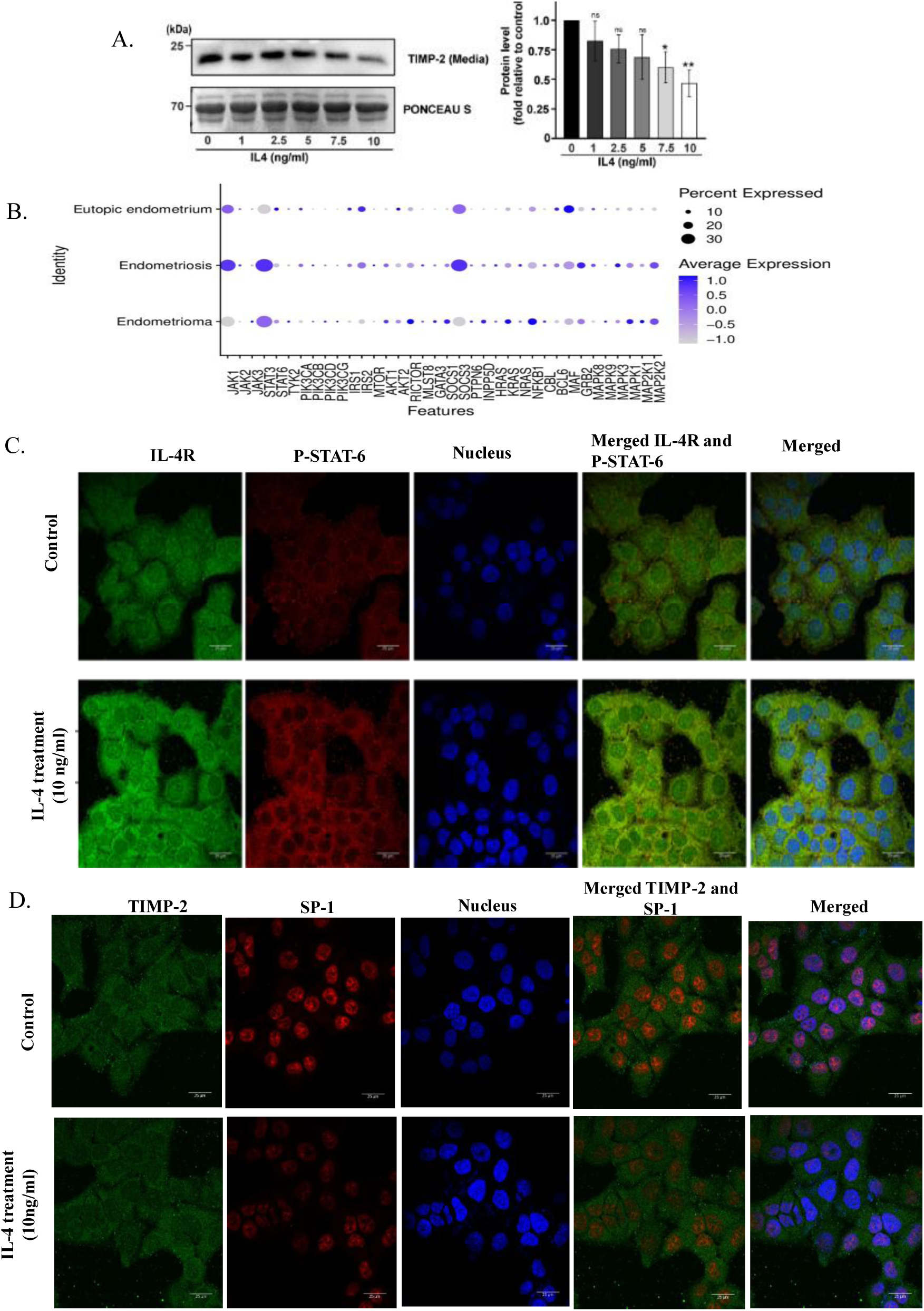
IL-4 suppresses TIMP-2 expression through SP1 modulation and activation of IL-4R/STAT6 signaling. (A) Dose-dependent effect of recombinant IL-4 (0, 1, 2.5, 5, 7.5, and 10 ng/ml) on TIMP-2 protein expression in Ishikawa cells, analyzed by immunoblotting. Corresponding densitometric quantification normalized to total protein loading (Ponceau S) is shown. (B) Single-cell transcriptomic analysis showing expression of genes associated with JAK/STAT, PI3K/AKT/MTOR, KRAS, and MAPK signaling pathways across eutopic endometrium, endometriosis, and endometrioma conditions. (C) Immunofluorescence analysis of IL-4R (green) and phosphorylated STAT6 (red) expression in untreated and IL-4-treated Ishikawa cells. Nuclei were counterstained with DAPI (blue). Merged images demonstrate activation of canonical IL-4 signaling following cytokine treatment. (D) Immunofluorescence analysis of TIMP-2 (green) and SP1 (red) expression in untreated and IL-4-treated Ishikawa cells. Nuclei were stained with DAPI (blue). IL-4 treatment resulted in reduced TIMP-2 and SP1 expression compared with untreated controls.

To further explore the molecular pathways that may mediate this effect, the expression of IL-4-associated signaling molecules was examined using the single-cell transcriptomic dataset. Components of the canonical IL-4 signaling pathway, including JAK1, JAK3, STAT6, together with downstream signaling molecules associated with PI3K, MAPK and transcriptional regulation, were detected across disease tissues, supporting the presence of an active IL-4 signaling network within the endometriotic microenvironment (Fig. 3B).

To experimentally validate activation of this pathway, immunofluorescence analysis was performed following IL-4 stimulation. Compared with untreated cells, IL-4-treated Ishikawa cells exhibited increased IL-4 receptor (IL-4R) expression accompanied by enhanced phosphorylation of STAT6, as demonstrated by stronger IL-4R and phospho-STAT6 immunoreactivity and increased co-localization in merged images (Fig. 3C). These findings indicate activation of the canonical IL-4R/STAT6 signaling pathway following IL-4 stimulation.

Since SP1 is a known transcriptional regulator of TIMP2, we next examined whether IL-4 signaling influences SP1 expression. Immunofluorescence analysis demonstrated that IL-4 treatment markedly reduced both TIMP-2 and SP1 expression compared with untreated cells (Fig. 3D). The concomitant reduction of SP1 and TIMP-2 following IL-4 stimulation suggests that IL-4-mediated suppression of TIMP-2 is associated with modulation of SP1-dependent transcription.

Collectively, these findings demonstrate that IL-4 suppresses TIMP-2 expression while activating the IL-4R/STAT6 signaling pathway. The simultaneous reduction of SP1 expression further suggests that SP1 may act downstream of IL-4 signaling to regulate TIMP-2 transcription, providing the basis for subsequent experiments investigating the transcriptional regulation of the TIMP-2 promoter.

### 3.4. SP1 positively regulates TIMP-2 expression and mediates IL-4-dependent transcriptional repression

Having demonstrated that IL-4 suppresses TIMP-2 expression and activates IL-4R/STAT6 signaling (Fig. 3), we next investigated whether the transcription factor SP1 mediates the inhibitory effect of IL-4 on TIMP-2 expression. Ishikawa cells were treated with increasing concentrations of recombinant IL-4 (0–10 ng/ml) for 12 h, followed by immunoblot analysis of SP1 and TIMP-2. IL-4 treatment resulted in a dose-dependent reduction in both SP1 and TIMP-2 protein expression, with significant suppression observed at 7.5 and 10 ng/ml (Fig. 4A), suggesting that IL-4-mediated downregulation of TIMP-2 is associated with reduced SP1 expression.

**Figure 4.**
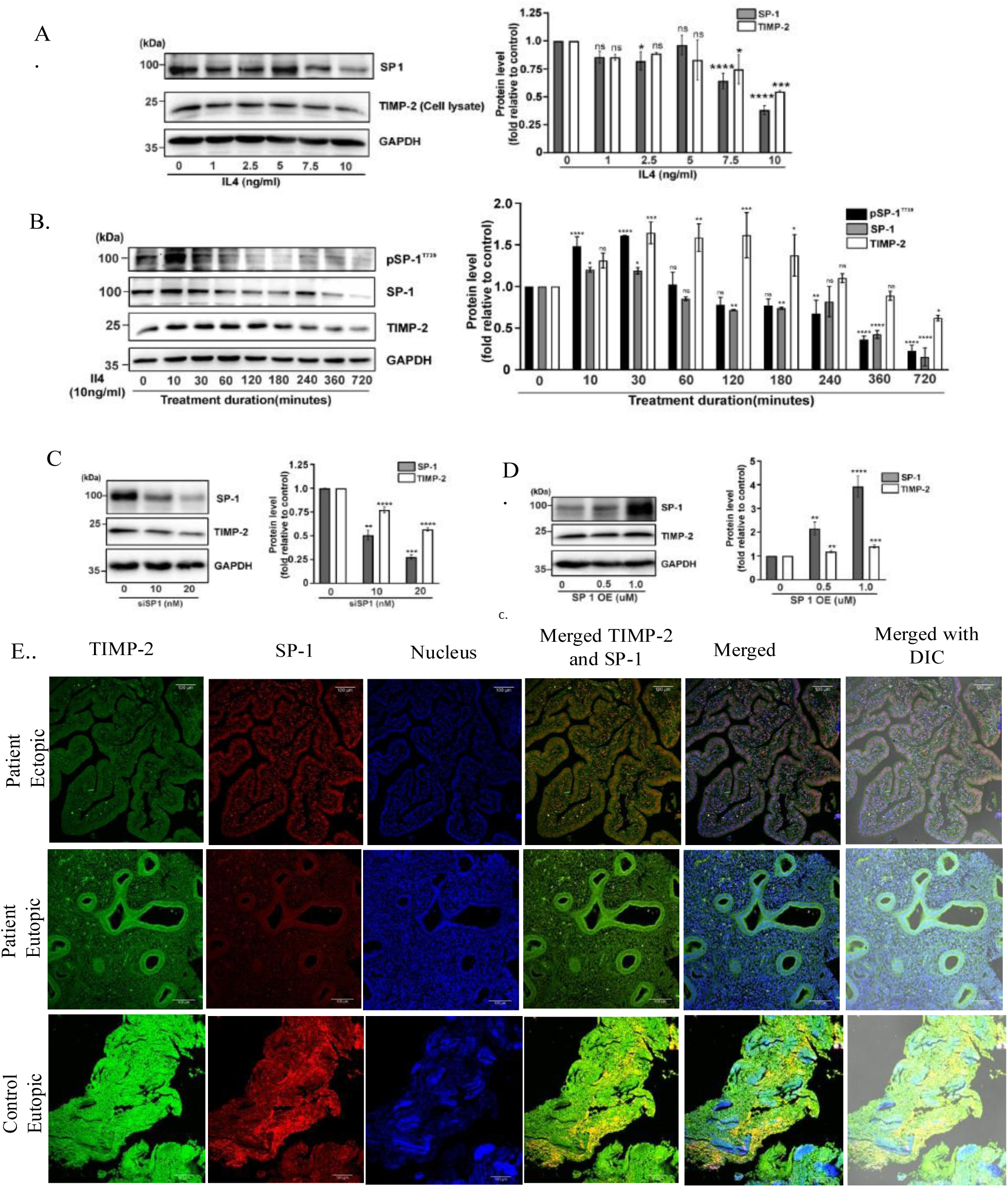
SP1 positively regulates TIMP-2 expression and mediates IL-4-dependent transcriptional repression. (A) Dose-dependent effect of recombinant IL-4 (0, 1, 2.5, 5, 7.5, and 10 ng/ml) on SP1 and TIMP-2 protein expression in Ishikawa cells, analyzed by immunoblotting. Corresponding densitometric analyses of SP1 and TIMP-2 normalized to GAPDH are shown. (B) Time-course analysis of IL-4 (10 ng/ml) treatment showing dynamic changes in phosphorylated SP1 (P-SP1), total SP1, and TIMP-2 protein levels. Bar graphs represent normalized protein expression relative to GAPDH. (C) siRNA-mediated SP1 knockdown (10 and 20 nM) results in reduced TIMP-2 expression, demonstrating the requirement of SP1 for TIMP-2 transcriptional regulation. (D) SP1 overexpression (0.5 and 1.0 µg plasmid DNA) enhances TIMP-2 protein expression in a dose-dependent manner. (E) Representative immunofluorescence images showing TIMP-2 (green) and SP1 (red) expression in control eutopic endometrium, eutopic endometrium from endometriosis patients, and ectopic endometriotic lesions. Nuclei were counterstained with DAPI (blue). Merged and DIC overlay images demonstrate co-localization and tissue morphology.

To determine the temporal relationship between SP1 activation and TIMP-2 expression, cells were treated with 10 ng/ml IL-4 for the indicated time intervals. Immunoblot analysis revealed a rapid increase in phosphorylated SP1 during the early phase of IL-4 stimulation, followed by a progressive decline in total SP1 expression with prolonged treatment. Consistent with this pattern, TIMP-2 expression gradually decreased over time, becoming markedly reduced at later time points (Fig. 4B). These findings indicate that IL-4 induces dynamic modulation of SP1 that precedes the suppression of TIMP-2 expression.

To directly examine the role of SP1 in regulating TIMP-2 expression, SP1 was transiently silenced using siRNA. Knockdown of SP1 resulted in a dose-dependent reduction in TIMP-2 protein expression compared with control cells (Fig. 4C), indicating that SP1 is required for maintaining basal TIMP-2 expression. Conversely, ectopic overexpression of SP1 significantly increased TIMP-2 protein levels in a dose-dependent manner (Fig. 4D), demonstrating that SP1 positively regulates TIMP-2 expression.

To validate these observations in clinical specimens, dual immunofluorescence staining was performed for TIMP-2 and SP1 in endometrial tissues. TIMP-2 expression was highest in control eutopic endometrium, reduced in eutopic endometrium from women with endometriosis, and further decreased in ectopic lesions. SP1 showed a similar reduction in eutopic endometrium from patients compared with controls (Fig. 4E). The concordant reduction of SP1 and TIMP-2 in patient tissues further supports the positive association between SP1 expression and TIMP-2 levels observed in vitro.

Collectively, these findings establish SP1 as a positive transcriptional regulator of TIMP-2 and demonstrate that IL-4 suppresses TIMP-2 expression through modulation of SP1. These observations prompted us to investigate whether genetic variation within the SP1-binding region of the TIMP-2 promoter influences IL-4-mediated transcriptional regulation.

### 3.5. Functional characterization of the TIMP2 −418 G/C promoter polymorphism reveals allele-specific transcriptional regulation by SP1 and IL-4

Having established that IL-4 suppresses TIMP-2 expression through modulation of SP1, we next investigated whether the functional −418 G/C polymorphism within the TIMP2 promoter influences SP1-dependent transcriptional regulation. GC+CC genotypes were significantly less frequent in patients than controls, consistent with a protective association of the C-containing genotypes in this cohort(Table 1.). Genotyping of clinical samples identified three genotypes (GG, GC and CC) among the study population (Fig. 5A). Western blot analysis further demonstrated that individuals carrying the CC genotype exhibited the highest TIMP-2 protein expression, whereas the GG genotype showed the lowest expression, with heterozygous GC samples displaying intermediate levels (Fig. 5B).

**Table 1.** Genotypic distribution of TIMP-2 -418G/C SNP in endometriosis in the eastern Indian population.

| Genotype | Patients<br>n= 117 | Control<br>n= 117 | OR | 95% CI | P value |
| --- | --- | --- | --- | --- | --- |
| GG | 94 | 82 | 1 |  |  |
| GC | 22 | 32 | 0.599 | 0.32- 1.113 | 0.07 |
| CC | 1 | 3 | 0.29 | 0.29-2.85 | 0.26 |
| GC+CC | 23 | 35 | 0.57 | 0.313-1.04 | **0.047** |

**Figure 5.**
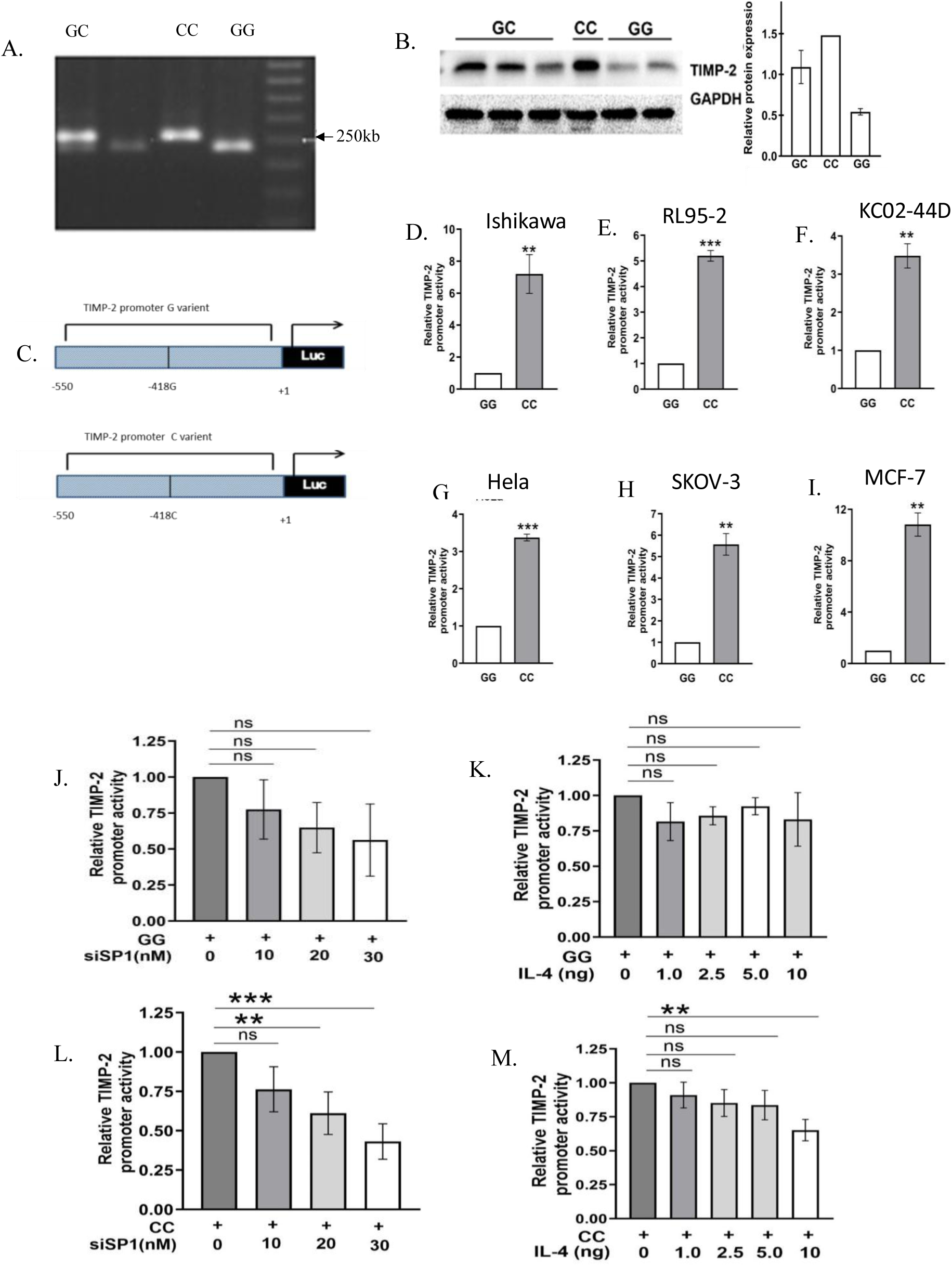
Functional characterization of the TIMP-2 −418G/C promoter polymorphism reveals allele-specific transcriptional regulation by SP1 and IL-4. (A) Representative PCR–RFLP analysis of the TIMP-2 −418G/C polymorphism showing GG, GC, and CC genotypes following BsoBI digestion. (B) Immunoblot analysis of TIMP-2 expression in ectopic endometrial tissues obtained from individuals carrying GG, GC, and CC genotypes. (C) Schematic representation of TIMP-2 promoter–luciferase reporter constructs containing either the −418G or −418C allele cloned into the pGL4.72 vector. (D–I) Basal promoter activities of G-and C-allele TIMP-2 constructs following transient transfection into Ishikawa, RL95-2, KC02-44D, HeLa, SKOV-3, and MCF-7 cells, respectively. Luciferase activity was normalized and expressed relative to the G allele construct. (J,L) Effect of SP1 siRNA-mediated knockdown (10, 20, and 30 nM) on TIMP-2 promoter activity. The C allele exhibited greater sensitivity to SP1 depletion compared with the G allele. (K,M) Effect of recombinant IL-4 treatment (0, 1, 2.5, 5, and 10 ng/ml) on TIMP-2 promoter activity. The −418C allele showed significantly greater repression in response to IL-4 compared with the G allele. Data are presented as mean ± SEM from three independent experiments. Statistical significance was determined using one-way ANOVA followed by Tukey’s post hoc test

**Figure 6.**
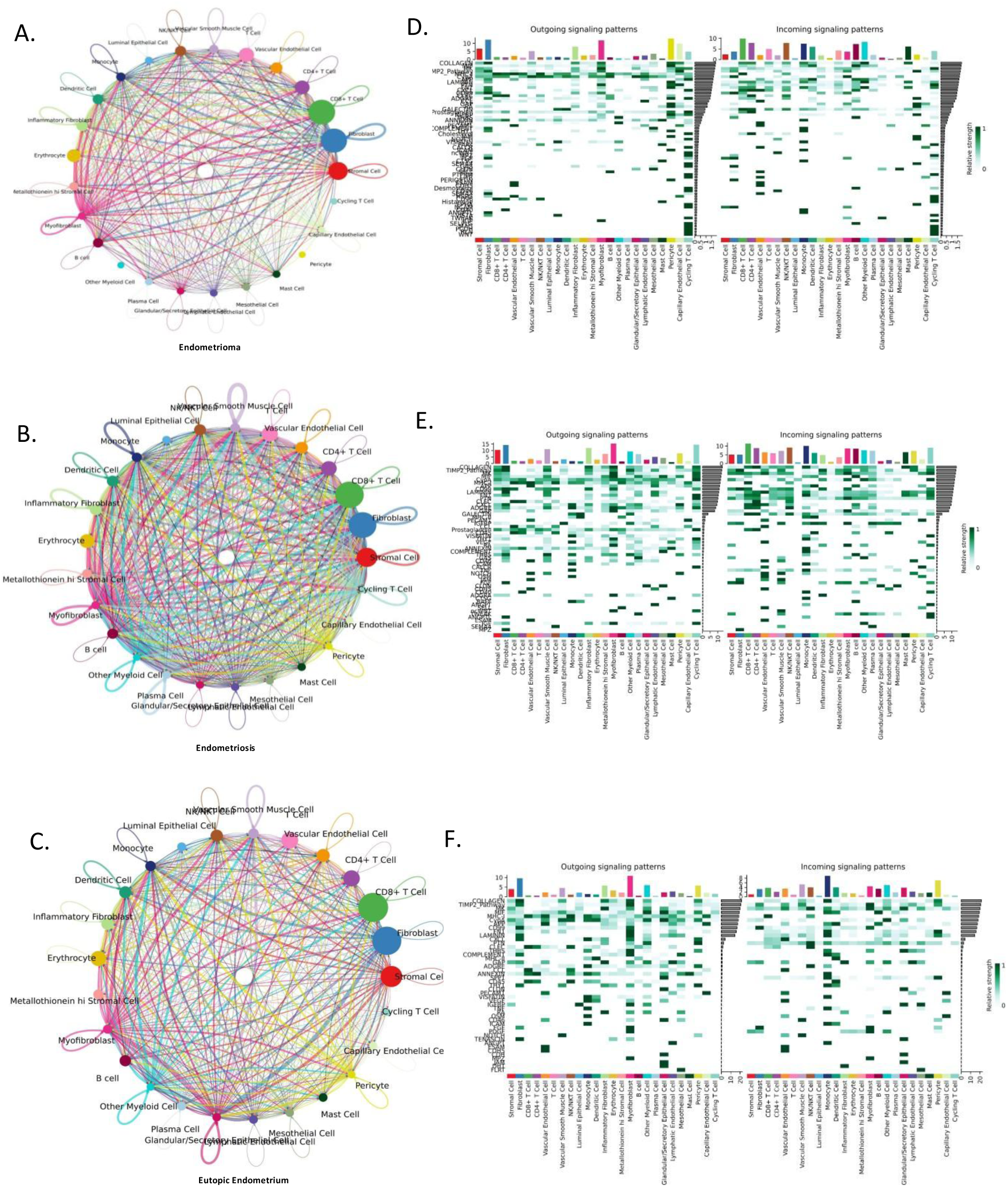
Cell–cell communication analysis identifies TIMP-2-associated signaling networks within the endometrial microenvironment. (A–C) Circle plots depicting cell–cell communication networks among distinct cellular populations in (A) endometrioma, (B) endometriosis, and (C) eutopic endometrium. Each node represents a specific cell type, while connecting lines indicate predicted intercellular interactions. The thickness of the connecting lines reflects the relative interaction strength between cellular populations. (D–F) Heatmaps showing pathway-specific outgoing and incoming signaling activities across different cell populations in (D) endometrioma, (E) endometriosis, and (F) eutopic endometrium. Rows represent signaling pathways and columns represent individual cell populations. Color intensity corresponds to the relative signaling activity, whereas bar plots above each heatmap indicate the overall contribution of each cell type to outgoing and incoming communication networks. Comparative analysis revealed altered intercellular communication patterns and extracellular matrix–associated signaling within the endometriotic microenvironment.

To determine whether this polymorphism directly affects promoter activity, approximately 550 bp promoter fragments containing either the −418G or −418C allele were cloned upstream of the luciferase reporter gene (Fig. 5C). Luciferase reporter assays performed in Ishikawa, RL95-2, KC02-44D, HeLa, SKOV-3 and MCF-7 cells consistently demonstrated significantly higher promoter activity for the −418C construct compared with the −418G construct (Fig. 5D–I), indicating that the C allele possesses greater transcriptional activity across multiple epithelial cell types.

Since SP1 positively regulated TIMP-2 expression (Fig. 4), we next examined whether promoter activity of the two alleles differed in response to SP1 depletion. Knockdown of SP1 produced only a modest, non-significant reduction in the activity of the −418G promoter (Fig. 5J). In contrast, SP1 silencing resulted in a significant dose-dependent decrease in −418C promoter activity (Fig. 5L), indicating that transcription driven by the C allele is more dependent on SP1.

Finally, we investigated whether the two promoter variants exhibit differential sensitivity to IL-4-mediated repression. Treatment with recombinant IL-4 produced minimal changes in −418G promoter activity (Fig. 5K), whereas the −418C promoter showed a progressive reduction in transcriptional activity, with significant inhibition observed at higher IL-4 concentrations (Fig. 5M). These findings indicate that the C allele is more responsive to IL-4-mediated transcriptional repression than the G allele.

Collectively, these results demonstrate that the −418 G/C polymorphism functionally regulates TIMP2 promoter activity in an allele-specific manner. The C allele exhibits greater basal transcriptional activity but is also more dependent on SP1 and more susceptible to IL-4-mediated repression, providing a molecular explanation for genotype-dependent regulation of TIMP2 expression in endometriosis.

#### TIMP-2 -418 G/C SNP and risk of Endometriosis

Odds Ratio (OR) was calculated by a binary logistic model by using Graph Pad In Stat. The p-value for the fisher’s exact test indicates the significance of the difference in genotype distributions between patients and controls. Significantly different values are shown in bold. CI-Confidence Interval, Ref = Reference genotype or allele for calculation of OR

### 3.6. Cell–cell communication analysis identifies TIMP2-associated signaling networks within the endometrial microenvironment

To understand intercellular communication within the endometrial microenvironment, we analyzed ligand–receptor interactions across eutopic endometrium, endometriosis, and endometrioma. The results revealed that endometriosis exhibited the highest number and strength of cell–cell interactions, followed by endometrioma and eutopic endometrium (Figure 7A–C). Immune cells such as cycling T cells, CD4⁺ and CD8⁺ T cells, monocytes, and dendritic cells displayed enhanced communication with mesenchymal cells including fibroblasts, stromal, and vascular smooth muscle cells, suggesting an active immune–stromal crosstalk in disease conditions. In contrast, epithelial cell–driven interactions predominated in endometrioma, reflecting its cystic and fibrotic nature.

We next examined the TIMP-2 signaling pathway, consisting of TIMP-2 as a ligand and LRP1, IGF1R, ITGA3, and MMP14 as receptors. This pathway showed stronger incoming and outgoing signaling activity in endometriosis compared to endometrioma and eutopic endometrium (Figure 7D–F). The enrichment was primarily observed in fibroblasts, myofibroblasts, inflammatory fibroblasts, stromal cells, and vascular smooth muscle cells, indicating that mesenchymal populations are key mediators of TIMP-2–related intercellular communication.

Together, these findings suggest that enhanced TIMP-2 signaling in mesenchymal cell clusters contributes to aberrant extracellular matrix remodeling and chronic inflammation in endometriosis. This context-specific activation of the TIMP-2 axis likely supports the invasive and persistent phenotype of endometrial lesions through coordinated stromal–immune interactions.

## 4. Discussion

TIMP-2 exerts both MMP-dependent and MMP-independent functions [36]; [37]. In its canonical MMP-dependent role, TIMP-2 regulates extracellular matrix proteolysis through inhibition of MMP activity and modulation of the MMP14 (MT1-MMP)/MMP-2 axis [38]; [14]. In contrast, its MMP-independent, ligand-like functions involve interactions with cell-surface proteins and receptors, including LRP1, IGF1R, and integrin α3 (ITGA3), through which TIMP-2 can modulate downstream signaling pathways governing cellular proliferation, apoptosis, migration, and angiogenic responses [37]; [39]; [40]. In our study cohort we observed significantly reduced TIMP-2 protein expression in ectopic lesions, eutopic endometrium, and serum samples from women with endometriosis.

Single-cell transcriptomic analysis showed that *TIMP2* was predominantly expressed in mesenchymal populations, particularly fibroblasts, myofibroblasts, stromal cells, and vascular smooth muscle cells. Fibroblasts exhibited both a high proportion of *TIMP2*-expressing cells and elevated average expression in ectopic lesions, suggesting that they are a major source of TIMP2 in the endometriotic microenvironment and may contribute to stromal ECM remodeling. Interestingly, myofibroblasts showed the highest average *TIMP2* expression in eutopic endometrium; given their established role in wound healing and fibrosis, and the recognized presence of fibrosis in the eutopic endometrium of women with endometriosis, this finding suggests that TIMP-2 may contribute to myofibroblast-mediated ECM remodeling [41]; [37]and fibrotic responses under eutopic conditions The reduced *TIMP2* expression in ectopic lesions may therefore reflect disruption of this homeostatic mechanism, leading to abnormal matrix remodeling during disease progression.

Although reduced TIMP-2 expression was consistently observed in endometriosis, the upstream inflammatory signals responsible for its dysregulation remained unclear. Endometriosis is initially characterized by a pro-inflammatory milieu; however, as lesions become established, the immune microenvironment gradually shifts towards a Th2-dominant, anti-inflammatory phenotype, which facilitates immune tolerance, fibrosis [42], extracellular matrix remodeling, and lesion persistence [43]; [44]. Among the Th2-associated cytokines, IL-4 is considered a key mediator of this chronic anti-inflammatory environment [43]. Consistent with this concept, our single-cell transcriptomic analysis also demonstrated increased Th2-associated immune signatures in ectopic lesions. These observations prompted us to investigate whether IL-4 contributes to TIMP-2 dysregulation during disease progression.

To further explore this possibility, we examined the cytokine profile across different tissue states using single-cell transcriptomic data. *IL4* emerged as one of the prominent cytokines enriched in ectopic lesions, while its receptor components, *IL4R* and *IL2RG*, also showed markedly elevated expression, indicating activation of IL-4 signaling within the endometriotic microenvironment. Functional validation using recombinant IL-4 demonstrated a dose-dependent reduction in TIMP2 secretion from endometrial epithelial cells. In parallel, transcriptomic analysis revealed increased expression of downstream IL-4 signaling molecules in ectopic tissues, suggesting activation of the canonical IL-4/STAT-6 pathway [45]; [46]. Consistent with these findings, IL-4 treatment markedly increased IL-4R and phosphorylated STAT6 levels while simultaneously reducing TIMP-2 expression at 10 ng/ml, supporting a role for IL-4 signaling in suppressing TIMP-2.

Although these findings established IL-4 as a negative regulator of TIMP2, the transcriptional mechanism underlying this regulation remained unknown. Previous characterization of the human TIMP2 promoter demonstrated the presence of five consensus SP1-binding sites [47], including one encompassing the −418 bp region, suggesting that SP1 is a major regulator of TIMP2 transcription [21]. Furthermore, previous studies have reported altered SP1 expression in endometriosis, making it a biologically relevant candidate. Based on these observations, we hypothesized that IL-4 suppresses TIMP2 through an SP1-dependent mechanism. We observed that IL-4 treatment reduced SP1 expression, whereas SP1 knockdown significantly decreased TIMP2 levels and SP1 overexpression restored TIMP2 expression, demonstrating that SP1 functions as a positive regulator of TIMP2 transcription under inflammatory conditions.

Immunofluorescence analysis further validated our in vitro observations at the tissue level. As expected, control eutopic endometrium exhibited the strongest TIMP2 and SP1 expression, whereas both proteins were reduced in tissues obtained from women with endometriosis. TIMP2 expression was lowest in ectopic lesions, consistent with our biochemical and transcriptomic findings. Interestingly, although SP1 staining appeared relatively higher in ectopic lesions than in matched eutopic endometrium from patients, this increase was not accompanied by restoration of TIMP2 expression. This observation suggests that SP1 abundance alone may not be sufficient to maintain TIMP2 transcription within the complex inflammatory microenvironment of ectopic lesions, where multiple regulatory pathways are likely to influence TIMP2 expression.

Furthermore, SP1 knockdown significantly reduced the promoter activity of the C allele construct, whereas the G allele construct showed no significant change. Similarly, IL-4 treatment selectively suppressed the promoter activity of the C allele without significantly affecting the G allele. Together, these findings suggest that the −418 G/C polymorphism differentially influences TIMP2 promoter activity and that the C allele is preferentially regulated by SP1 and IL-4-mediated inflammatory signaling.

Functional promoter polymorphisms are increasingly recognized as important determinants of disease susceptibility because they can influence transcription factor binding and gene expression. The TIMP2 −418 G/C (rs8179090) promoter polymorphism has been investigated in women with endometriosis from North Chinese and Korean populations [20], [18], as well as in abdominal aortic aneurysm [19], moyamoya disease [48], age-related macular degeneration [49], and cervical cancer [50], although the reported associations have been inconsistent across different populations. However, the functional significance of rs8179090 in regulating TIMP2 expression during chronic inflammatory conditions has remained largely unexplored.

We next investigated whether the TIMP2 −418 G/C (rs8179090) promoter polymorphism influences TIMP2 transcription. In the present study, genotype analysis revealed that the combined GC+CC genotypes were significantly less frequent among women with endometriosis than healthy controls (OR = 0.57, *P* = 0.047), suggesting that the C allele may confer protection against disease susceptibility, whereas the GG genotype was more prevalent among affected individuals. Although the GC genotype alone showed a similar trend toward reduced risk, it did not reach statistical significance, and the CC genotype was too infrequent to allow meaningful independent analysis. Patients carrying C-containing genotypes/allele are associated with higher TIMP2 expression and higher promoter activity, while CC is rare in the cohort suggesting that this promoter variant contributes to inter-individual differences in TIMP2 expression. To validate the functional significance of this polymorphism, we generated promoter-reporter constructs containing either the G or C allele and examined their transcriptional activity in epithelial cell lines. Consistent with the patient data, the C allele displayed markedly higher basal promoter activity than the G allele, indicating that the −418 G/C polymorphism functionally regulates TIMP2 transcription. Previous studies have proposed that rs8179090 is located within a consensus SP1-binding motif of the TIMP2 promoter and may therefore influence SP1-dependent transcription. Although these studies were based primarily on sequence prediction, our findings provide functional evidence that promoter activity differs substantially between the two alleles. Furthermore, modulation of SP1 expression by siRNA-mediated knockdown or SP1 overexpression, together with IL-4 treatment, supports the concept that SP1 is a positive regulator of TIMP2 transcription and that inflammatory signaling influences this promoter in an allele-dependent manner.

To explore the biological significance of reduced TIMP2 expression, we analyzed TIMP2-mediated cell–cell communication within the endometrial microenvironment. CellChat analysis revealed enhanced intercellular communication in endometriosis and endometrioma compared with eutopic endometrium, with TIMP2 acting as the signaling ligand and LRP1, IGF1R, ITGA3, and MMP14 serving as the major receptor components [37], predominantly among mesenchymal cell populations. These findings suggest that dysregulation of TIMP2-mediated signaling may contribute not only to extracellular matrix remodeling but also to altered stromal–immune communication that supports lesion persistence in endometriosis.

To our knowledge, this is the first study to investigate the functional significance of the TIMP2 −418 G/C (rs8179090) promoter polymorphism in endometriosis in an Eastern Indian cohort. Our findings demonstrate that IL-4 suppresses TIMP2 expression through SP1-dependent transcriptional regulation and reveal that the C allele exhibits higher basal promoter activity and greater responsiveness to SP1 depletion and IL-4-mediated repression than the G allele. Together with the altered TIMP2-mediated cell–cell communication observed in endometriotic tissues, these findings support a model in which inflammatory signaling and promoter variation coordinately regulate TIMP2 expression during disease progression. Further studies investigating allele-specific SP1 occupancy and in vivo regulation of the TIMP2 promoter will help clarify the molecular basis of this regulatory pathway and its therapeutic potential in endometriosis.

## Author contributions

Susmita Saha: Conceptualization (supporting); Formal Analysis; Investigation; Methodology; Validation; Visualization; Writing— original draft, review, and editing. Supratim Ghosh: Single-cell data curation, Formal analysis and interpretation; Writing—review editing (supporting). Tapasi Roy: Writing—review and editing. Debasis Nayak: Resources. Pramathes Das Mahapatra: Resources; Supervision (supporting). Arindam Maitra: Supervision (supporting). Snehasikta Swarnakar: Conceptualization (lead); Supervision (lead); Writing— review, and editing (supporting).

## Acknowledgments

Susmita Saha is thankful to the University Grants Commission (UGC), India, for providing financial support. We gratefully acknowledge Dr. Geetanjali Sachdeva for providing the Ishikawa cell line and Dr. Deepak Modi for providing the RL95-2 and KC02-44D hTERT cell lines. We also acknowledge Director, CSIR-Indian Institute of Chemical Biology (CSIR-IICB), Kolkata, India, for her kind support.

## References

[1] B. Smolarz, K. Szyllo, H. Romanowicz, Endometriosis: Epidemiology, Classification, Pathogenesis, Treatment and Genetics (Review of Literature), Int J Mol Sci, 22 (2021).

[2] H. Mariadas, J.H. Chen, K.H. Chen, The Molecular and Cellular Mechanisms of Endometriosis: From Basic Pathophysiology to Clinical Implications, Int J Mol Sci, 26 (2025).

[3] Y. Wang, K. Nicholes, I.M. Shih, The Origin and Pathogenesis of Endometriosis, Annu Rev Pathol, 15 (2020) 71−95.

[4] R.J. Consoli, K. Carlson, Endometriosis, StatPearls, Place Published, 2026.

[5] E. Somigliana, P. Vigano, L. Benaglia, A. Busnelli, P. Vercellini, L. Fedele, Adhesion prevention in endometriosis: a neglected critical challenge, J Minim Invasive Gynecol, 19 (2012) 415−421.

[6] S.E. Elizur, J. Mostafa, E. Berkowitz, R. Orvieto, Endometriosis and infertility: pathophysiology, treatment strategies, and reproductive outcomes, Arch Gynecol Obstet, 312 (2025) 1037−1048.

[7] S. Matsuzaki, C. Darcha, J.L. Pouly, M. Canis, Effects of matrix stiffness on epithelial to mesenchymal transition−like processes of endometrial epithelial cells: Implications for the pathogenesis of endometriosis, Sci Rep, 7 (2017) 44616.

[8] G. Vissers, M. Giacomozzi, W. Verdurmen, R. Peek, A. Nap, The role of fibrosis in endometriosis: a systematic review, Hum Reprod Update, 30 (2024) 706−750.

[9] J.M. Garcia Garcia, V. Vannuzzi, C. Donati, C. Bernacchioni, P. Bruni, F. Petraglia, Endometriosis: Cellular and Molecular Mechanisms Leading to Fibrosis, Reprod Sci, 30 (2023) 1453−1461.

[10] R. Sadat Sandoghsaz, F. Montazeri, H. Shafienia, S. Mehdi Kalantar, A. Javaheri, M. Samadi, Expression of miR−21 &IL−4 in endometriosis, Hum Immunol, 85 (2024) 110746.

[11] A. Drosdzol−Cop, V. Skrzypulec−Plinta, R. Stojko, Serum and peritoneal fluid immunological markers in adolescent girls with chronic pelvic pain, Obstet Gynecol Surv, 67 (2012) 374−381.

[12] A.M. Malutan, T. Drugan, R. Ciortea, R.F. Mocan−Hognogi, C. Bucuri, M.P. Rada, D. Mihu, Serum anti−inflammatory cytokines for the evaluation of inflammatory status in endometriosis, J Res Med Sci, 20 (2015) 668−674.

[13] Y.Y. Fan, H.Y. Chen, W. Chen, Y.N. Liu, Y. Fu, L.N. Wang, Expression of inflammatory cytokines in serum and peritoneal fluid from patients with different stages of endometriosis, Gynecol Endocrinol, 34 (2018) 507−512.

[14] M.M. Bernardo, R. Fridman, TIMP−2 (tissue inhibitor of metalloproteinase−2) regulates MMP−2 (matrix metalloproteinase−2) activity in the extracellular environment after pro−MMP−2 activation by MT1 (membrane type 1)−MMP, Biochem J, 374 (2003) 739−745.

[15] K. Brew, H. Nagase, The tissue inhibitors of metalloproteinases (TIMPs): an ancient family with structural and functional diversity, Biochim Biophys Acta, 1803 (2010) 55−71.

[16] S. Jana, K. Chatterjee, A.K. Ray, P. DasMahapatra, S. Swarnakar, Regulation of Matrix Metalloproteinase−2 Activity by COX−2−PGE2−pAKT Axis Promotes Angiogenesis in Endometriosis, PLoS One, 11 (2016) e0163540.

[17] T. Wu, R. Zhang, Q. Jiang, Z. Li, R. Wu, Expression of cellular adherent and invasive molecules in recurrent ovarian endometriosis, J Int Med Res, 48 (2020) 300060520971993.

[18] Y.J. Cho, N.H. Kim, K.A. Jeong, J.Y. Lee, H.S. Moon, H.L. Kim, H.W. Chung, Association between MMP−2 and TIMP−2 gene polymorphisms and advanced−stage endometriosis in Korean women, Am J Reprod Immunol, 69 (2013) 73−84.

[19] J. Mikolajczyk−Stecyna, A. Korcz, M. Gabriel, K. Pawlaczyk, G. Oszkinis, R. Slomski, Gene polymorphism −418 G/C of tissue inhibitor of metalloproteinases 2 is associated with abdominal aortic aneurysm, J Vasc Surg, 61 (2015) 1114−1119.

[20] S. Kang, X.W. Zhao, N. Wang, S.C. Chen, R.M. Zhou, Y. Li, Association of polymorphisms of the MMP−2 and TIMP−2 genes with the risk of endometriosis in North Chinese women, Fertil Steril, 90 (2008) 2023−2029.

[21] K. Hirano, T. Sakamoto, Y. Uchida, Y. Morishima, K. Masuyama, Y. Ishii, A. Nomura, M. Ohtsuka, K. Sekizawa, Tissue inhibitor of metalloproteinases−2 gene polymorphisms in chronic obstructive pulmonary disease, Eur Respir J, 18 (2001) 748−752.

[22] Y. Zhou, C. Yu, X. Miao, W. Tan, G. Liang, P. Xiong, T. Sun, D. Lin, Substantial reduction in risk of breast cancer associated with genetic polymorphisms in the promoters of the matrix metalloproteinase−2 and tissue inhibitor of metalloproteinase−2 genes, Carcinogenesis, 25 (2004) 399−404.

[23] Y.A. De Clerck, M.I. Darville, Y. Eeckhout, G.G. Rousseau, Characterization of the promoter of the gene encoding human tissue inhibitor of metalloproteinases−2 (TIMP−2), Gene, 139 (1994) 185−191.

[24] Z.D. Zhong, K. Hammani, W.S. Bae, Y.A. DeClerck, NF−Y and Sp1 cooperate for the transcriptional activation and cAMP response of human tissue inhibitor of metalloproteinases−2, J Biol Chem, 275 (2000) 18602−18610.

[25] A.M. Malutan, C. Drugan, T. Drugan, R. Ciortea, D. Mihu, The association between interleukin−4 − 590C/T genetic polymorphism, IL−4 serum level, and advanced endometriosis, Cent Eur J Immunol, 41 (2016) 176−181.

[26] K.K. Chang, L.B. Liu, L.P. Jin, B. Zhang, J. Mei, H. Li, C.Y. Wei, W.J. Zhou, X.Y. Zhu, J. Shao, D.J. Li, M.Q. Li, IL−27 triggers IL−10 production in Th17 cells via a c−Maf/RORgammat/Blimp−1 signal to promote the progression of endometriosis, Cell Death Dis, 8 (2017) e2666.

[27] W.J. Zhou, H.L. Yang, J. Shao, J. Mei, K.K. Chang, R. Zhu, M.Q. Li, Anti−inflammatory cytokines in endometriosis, Cell Mol Life Sci, 76 (2019) 2111−2132.

[28] A. Junga, M. Pilmane, Z. Abola, O. Volrats, The Morphopathogenetic Aspects of Intraabdominal Adhesions in Children under One Year of Age, Medicina (Kaunas), 55 (2019).

[29] M.A.S. Fonseca, M. Haro, K.N. Wright, X. Lin, F. Abbasi, J. Sun, L. Hernandez, N.L. Orr, J. Hong, Y. Choi−Kuaea, H.M. Maluf, B.L. Balzer, A. Fishburn, R. Hickey, I. Cass, H.S. Goodridge, M. Truong, Y. Wang, M.D. Pisarska, H.Q. Dinh, A. El−Naggar, D.G. Huntsman, M.S. Anglesio, M.T. Goodman, F. Medeiros, M. Siedhoff, K. Lawrenson, Single−cell transcriptomic analysis of endometriosis, Nat Genet, 55 (2023) 255−267.

[30] Y. Hao, S. Hao, E. Andersen−Nissen, W.M. Mauck, 3rd, S. Zheng, A. Butler, M.J. Lee, A.J. Wilk, C. Darby, M. Zager, P. Hoffman, M. Stoeckius, E. Papalexi, E.P. Mimitou, J. Jain, A. Srivastava, T. Stuart, L.M. Fleming, B. Yeung, A.J. Rogers, J.M. McElrath, C.A. Blish, R. Gottardo, P. Smibert, R. Satija, Integrated analysis of multimodal single−cell data, Cell, 184 (2021) 3573−3587 e3529.

[31] C.S. McGinnis, L.M. Murrow, Z.J. Gartner, DoubletFinder: Doublet Detection in Single−Cell RNA Sequencing Data Using Artificial Nearest Neighbors, Cell Syst, 8 (2019) 329−337 e324.

[32] C. Hafemeister, R. Satija, Normalization and variance stabilization of single−cell RNA−seq data using regularized negative binomial regression, Genome Biol, 20 (2019) 296.

[33] I. Korsunsky, N. Millard, J. Fan, K. Slowikowski, F. Zhang, K. Wei, Y. Baglaenko, M. Brenner, P.R. Loh, S. Raychaudhuri, Fast, sensitive and accurate integration of single−cell data with Harmony, Nat Methods, 16 (2019) 1289−1296.

[34] S. Jin, C.F. Guerrero−Juarez, L. Zhang, I. Chang, R. Ramos, C.H. Kuan, P. Myung, M.V. Plikus, Q. Nie, Inference and analysis of cell−cell communication using CellChat, Nat Commun, 12 (2021) 1088.

[35] R. Kapoor, C.A. Stratopoulou, M.M. Dolmans, Pathogenesis of Endometriosis: New Insights into Prospective Therapies, Int J Mol Sci, 22 (2021).

[36] V. Arpino, M. Brock, S.E. Gill, The role of TIMPs in regulation of extracellular matrix proteolysis, Matrix Biol, 44−46 (2015) 247−254.

[37] D. Peeney, Y. Liu, C. Lazaroff, S. Gurung, W.G. Stetler−Stevenson, Unravelling the distinct biological functions and potential therapeutic applications of TIMP2 in cancer, Carcinogenesis, 43 (2022) 405−418.

[38] H.W. Chung, J.Y. Lee, H.S. Moon, S.E. Hur, M.H. Park, Y. Wen, M.L. Polan, Matrix metalloproteinase− 2, membranous type 1 matrix metalloproteinase, and tissue inhibitor of metalloproteinase−2 expression in ectopic and eutopic endometrium, Fertil Steril, 78 (2002) 787−795.

[39] H. Emonard, G. Bellon, L. Troeberg, A. Berton, A. Robinet, P. Henriet, E. Marbaix, K. Kirkegaard, L. Patthy, Y. Eeckhout, H. Nagase, W. Hornebeck, P.J. Courtoy, Low density lipoprotein receptor−related protein mediates endocytic clearance of pro−MMP−2.TIMP−2 complex through a thrombospondin− independent mechanism, J Biol Chem, 279 (2004) 54944−54951.

[40] D.W. Seo, H. Li, L. Guedez, P.T. Wingfield, T. Diaz, R. Salloum, B.Y. Wei, W.G. Stetler−Stevenson, TIMP− 2 mediated inhibition of angiogenesis: an MMP−independent mechanism, Cell, 114 (2003) 171−180.

[41] L. Costanzo, B. Soto, R. Meier, P. Geraghty, The Biology and Function of Tissue Inhibitor of Metalloproteinase 2 in the Lungs, Pulm Med, 2022 (2022) 3632764.

[42] D. Feng, Y. Li, H. Zheng, Y. Wang, J. Deng, T. Liu, W. Liao, F. Shen, IL−4−induced M2 macrophages inhibit fibrosis of endometrial stromal cells, Reprod Biol, 24 (2024) 100852.

[43] K. Szymanowski, J. Niepsuj−Binias, A. Dera−Szymanowska, M. Wolun−Cholewa, A. Yantczenko, E. Florek, T. Opala, M. Murawski, K. Wiktorowicz, An influence of immunomodulation on Th1 and Th2 immune response in endometriosis in an animal model, Biomed Res Int, 2013 (2013) 849492.

[44] L. Krygere, P. Jukna, K. Jariene, E. Drejeriene, Diagnostic Potential of Cytokine Biomarkers in Endometriosis: Challenges and Insights, Biomedicines, 12 (2024).

[45] K. Takeda, T. Tanaka, W. Shi, M. Matsumoto, M. Minami, S. Kashiwamura, K. Nakanishi, N. Yoshida, T. Kishimoto, S. Akira, Essential role of Stat6 in IL−4 signalling, Nature, 380 (1996) 627−630.

[46] M. Pesu, K. Takaluoma, S. Aittomaki, A. Lagerstedt, K. Saksela, P.E. Kovanen, O. Silvennoinen, Interleukin−4−induced transcriptional activation by stat6 involves multiple serine/threonine kinase pathways and serine phosphorylation of stat6, Blood, 95 (2000) 494−502.

[47] K. Hammani, A. Blakis, D. Morsette, A.M. Bowcock, C. Schmutte, P. Henriet, Y.A. DeClerck, Structure and characterization of the human tissue inhibitor of metalloproteinases−2 gene, J Biol Chem, 271 (1996) 25498−25505.

[48] Y.S. Park, Y.J. Jeon, H.S. Kim, I.B. Han, S.H. Oh, D.S. Kim, N.K. Kim, The GC + CC genotype at position −418 in TIMP−2 promoter and the −1575GA/−1306CC genotype in MMP−2 is genetic predisposing factors for prevalence of moyamoya disease, BMC Neurol, 14 (2014) 180.

[49] H. Ortak, S. Demir, O. Ates, I. Benli, E. Sogut, M. Sahin, The role of MMP2 (−1306C>T) and TIMP2 (−418 G>C) promoter variants in age−related macular degeneration, Ophthalmic Genet, 34 (2013) 217−222.

[50] P. Srivastava, S. Pandey, B. Mittal, R.D. Mittal, No Association of Matrix Metalloproteinase [MMP]− 2 (−735C > T) and Tissue Inhibitor of Metalloproteinase [TIMP]−2 (−418G > C) Gene Polymorphisms with Cervical Cancer Susceptibility, Indian J Clin Biochem, 28 (2013) 13−18.

